# The differential contribution of pacemaker neurons to synaptic transmission in the pyloric network of the Jonah crab, *Cancer borealis*

**DOI:** 10.1101/523829

**Authors:** Diana Martinez, Joseph M. Santin, David Schulz, Farzan Nadim

## Abstract

Many neurons receive synchronous input from heterogeneous presynaptic neurons with distinct properties. An instructive example is the crustacean stomatogastric pyloric circuit pacemaker group, consisting of the anterior burster (AB) and pyloric dilator (PD) neurons, which are active synchronously and exert a combined synaptic action on most pyloric follower neurons. Although the stomatogastric system of the crab *Cancer borealis* has become a preferred model system for exploration of cellular and synaptic basis of circuit dynamics, in this species, the identity of the PD neuron neurotransmitter and its contribution to the total pacemaker group synaptic output remain unexplored. We examined the synaptic properties of the crab PD neuron using a combination of single cell mRNA analysis, electrophysiology and pharmacology. The crab PD neuron expresses high levels of choline acetyltransferase and the vesicular acetylcholine transporter mRNAs, hallmarks of cholinergic neurons. Conversely, the AB neuron does not express either of these cholinergic markers, and expresses high levels of vesicular glutamate transporter mRNA, consistent with a glutamatergic phenotype. Notably, in the combined synapses to the LP and PY neurons, the major contribution is from the glutamatergic AB neuron and only between 25-30% of the synaptic strength is due to the PD neuron. However, there was no difference between the short-term synaptic plasticity in the total pacemaker synapse compared to that of the PD neuron alone. These findings provide a guide for similar explorations of heterogeneous synaptic connections in other systems and a baseline in this system for the exploration of the differential influence of neuromodulators.

## New and Noteworthy

The pacemaker-driven pyloric circuit of the Jonah crab stomatogastric nervous system is a well-studied model system for exploring circuit dynamics and neuromodulation. Yet the understanding of the synaptic properties of the two pacemaker neuron types is based on older analyses in other species. We use single-cell PCR and electrophysiology to explore the neurotransmitters the pacemaker neurons and their distinct contribution to the combined synaptic potentials.

## Introduction

Within neural circuits, the activity pattern of any neuron is shaped by input from multiple presynaptic neurons. Inputs from distinct presynaptic neurons may have different strengths or signs, and may be subject to different modulatory modifications (Johnson et al. 2011). Even when multiple presynaptic neurons are active synchronously, differences in the strength, dynamics and plasticity rules of the synapses modify the total influence on the postsynaptic neuron (Abbott et al. 1997; Markram et al. 1998; Puccini et al. 2007). These differences depend on neurotransmitter types, presence of co-transmitters, and on the network activity and modulatory state. Consequently, properties of distinct presynaptic inputs are important factors in defining the neural circuit output and its plasticity in response to different brain states and behavioral needs (Nadim and Bucher 2014).

The pyloric circuit of the crustacean stomatogastric ganglion provides a noteworthy example of flexibility resulting from multiple synaptic inputs (Daur et al. 2016). The pyloric circuit oscillatory activity is driven by a pacemaker group, consisting of one anterior burster (AB) and two pyloric dilator (PD) neurons. These neurons are connected through gap junction-mediated electrical coupling and produce synchronous bursting oscillations. Previous studies of this neural circuit in the spiny lobster *Panulirus interruptus* have shown that the AB and PD neurons have distinct neurotransmitter types: the AB neuron is glutamatergic, whereas the PD neurons are cholinergic (Eisen and Marder 1982; Marder and Eisen 1984b). Consequently, the postsynaptic influences of these two neuron types are distinct in their strength and kinetics (Rabbah and Nadim 2007). However, most postsynaptic targets of the pyloric pacemaker group, including the lateral pyloric (LP) and pyloric constrictor (PY) neurons, receive synaptic input from both AB and PD neurons. As a result, although these postsynaptic neurons experience a composite synaptic input from the pacemaker group, the relative contributions of the AB and PD synapses to this total current are distinct in their strength, kinetics and its response to modulatory inputs (Rabbah and Nadim 2007).

Most recent studies of the pyloric circuit use the crab *Cancer borealis* as the experimental species (Blitz 2017; Follmann et al. 2017; Haddad and Marder 2018; Haley et al. 2018; Lett et al. 2017; Li et al. 2018; Otopalik et al. 2017; White et al. 2017). In this species, however, the neurotransmitters used by the pacemaker AB and PD neurons have not been described, and the relative contribution of the AB and PD synapses remains unquantified. In this study, we examine the neurotransmitter identity of the AB and PD neurons in *C. borealis*, explore the contribution of the AB and PD synapses to the total synaptic input in the LP and PY neurons, and examine the short-term dynamics of these synapses.

## Methods

Adult male crabs (*Cancer borealis*) were acquired from local distributors and maintained in aquaria filled with chilled artificial saline until ready for use. Prior to dissection, crabs were placed on ice and the dissection was performed using standard protocols as previously described (Blitz and Nusbaum 2012; Tseng and Nadim 2010). The stomatogastric nervous system, including the four ganglia (esophageal ganglion, two commissural ganglia and the stomatogastric ganglion, STG), the motor nerves and connecting nerves, were dissected from the stomach and pinned to a saline filled, Sylgard (Dow-Corning) lined 100 mm Petri dish. The STG was desheathed, exposing the somata of the neurons for intracellular impalement. Preparations were superfused with chilled (11-13°C) physiological *Cancer* saline containing: 11 mM KCl, 440 mM NaCl, 13 mM CaCl_2_ · 2H_2_O, 26 mM MgCl_2_ · 6H_2_O, 11.2 mM Trizma base, 5.1 mM maleic acid, adjusted to a pH of 7.4.

Extracellular recordings were obtained from identified motor nerves using stainless steel electrodes, placed inside and outside of a petroleum jelly well created to electrically isolate a small section of the nerve, and amplified using a differential AC amplifier model 1700 (A-M Systems). Individual pyloric neurons were impaled with glass microelectrodes and identified via their activity patterns, axonal projections in recorded motor nerves and their synaptic interactions with other neurons within the network (Weimann et al. 1991). Intracellular glass microelectrodes were prepared using the Flaming-Brown micropipette puller (P97; Sutter Instruments) and filled with 0.6 M K_2_SO_4_ and 20 mM KCl. Current injections were done using microelectrodes with a resistance of 15-22 MΩ; membrane potential measurements were performed using microelectrodes with a resistance of 25-30 MΩ. Intracellular recordings were performed using Axoclamp 2B and 900A amplifiers (Molecular Devices) and digitized using the Digidata acquisition board and pClamp 9 software (Molecular Devices).

### Comparison of Synapses

The synapses between pyloric neurons have two components: spike mediated and graded (Graubard et al. 1980). In this study we focus on the graded component, the dominant mode of transmission in pyloric synapses (Manor et al. 1997; Rosenbaum and Marder 2018; Zhao et al. 2011), which we measured in the presence of 10^−7^ M tetrodotoxin (TTX; Biotium). TTX blocks sodium channels and therefore removes all action potential activity. In the presence of TTX, all pyloric neurons become quiescent at a resting potential of around −60 mV. TTX also blocks all descending modulatory input to the STG.

Our measurements of synaptic potentials include synaptic inputs from both AB and PD neurons. We controlled the membrane potential of the pacemaker ensemble by controlling the PD neuron in two-electrode voltage clamp (TEVC) mode (Li et al. 2018). We measured the postsynaptic potential (due to both pacemaker neuron types) simultaneously in the LP and PY neurons (Rabbah and Nadim 2007). The PD neuron was voltage clamped at −60mV. The voltage-clamped PD neuron was then depolarized with a train of five pulses (duration: 500 ms; inter-pulse interval: 500 ms) with amplitudes ranging from 15 to 45 mV, in 5 mV intervals. The postsynaptic neurons (PY and LP) were each held at −50 mV, away from the synaptic reversal potential of around −80mV, in order to record the synaptic current, using a single-electrode in discontinuous current clamp (DCC). Following measurement of the synapses in control saline (including 10^−7^ M TTX), to isolate the cholinergic component of the total postsynaptic potential, we repeated our measurements after 30 min superfusion in 10^−7^ M TTX and 10^−5^ M picrotoxin (PTX; Sigma-Aldrich). PTX blocks glutamatergic synapses in the STG of both *C. borealis* and *P. interruptus* (Bidaut 1980; Cleland and Selverston 1995; Marder and Paupardin-Tritsch 1978; Rinberg et al. 2013; Temporal et al. 2012) and, in the latter species has been found to selectively block synaptic inhibition from the AB neuron but not that from the PD neuron (Rabbah and Nadim 2007). After measurements in PTX, the preparations were superfused with control saline for 45 minutes for measurements in wash conditions.

To quantify the synaptic strength, we fit the difference between the postsynaptic baseline voltage and the peak amplitude of the postsynaptic potential, evaluated at all presynaptic voltages, with a Boltzmann sigmoidal equation:

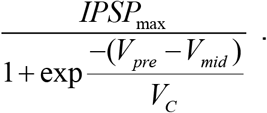

For each condition, we evaluated the maximum amplitude (IPSP_max_), activation midpoint value (V_mid_) and activation slope factor (V_C_).

Short-term synaptic dynamics were measured in both LP and PY neurons in response to a train of five 500 ms square pulses with 500 ms inter-pulse intervals (Rabbah and Nadim 2005). The extent of synaptic depression in both LP and PY neurons was quantified as a ratio of the response of the fifth pulse to the first pulse.

### Harvesting identified neurons

The PD, LP, PY, and AB neurons were identified as described above. Due to their small soma size, the AB neurons were filled with Alexa Fluor 488 at a concentration of 4 mg/ml (ThermoFisher Scientific) by electrophoresis for 20 mins with a constant −20 nA DC current, to ease identification for harvesting after electrophysiological identification. Cells then were collected as described previously (Schulz et al. 2007). Briefly, after electrophysiological identification of neurons, a petroleum jelly well was built around the STG. The ganglion was then exposed to ~2.5 mg/mL protease (Sigma Aldrich) diluted in crab saline to digest the connective tissue. After loosening of the neurons by protease digestion, the protease was thoroughly washed away from the ganglion and the saline in the well was replaced by 70% ethylene glycol diluted in *Cancer* saline over the course of ~15-20 min. The Petri dish was then placed at −20°C for 1 hour. A total of 5 PD, 5 LP, 6 PY, and 7 AB neurons were hand dissected from the STG (N=7 ganglia) using fine forceps. Each neuron was placed in 400 μL lysis buffer and stored at −80°C.

### cDNA synthesis and pre-amplification

Total RNA was isolated from each neuron using the Quick-RNA MicroPrep kit (Zymo Research) per manufacturer’s instructions. Reverse transcription of total RNA was then performed using a mixture of oligo-dT and random hexamer primers (qScript cDNA Supermix; QuantaBio). Half of the cDNA produced from each neuron (10 μL) was then pre-amplified using PerfeCTa PreAmp Supermix (QuantaBio) according to the manufacturer’s instructions (20 μL reaction volume) with 0.5 μM final concentration of each gene-specific primer (Table 1). This protocol utilizes a 14-cycle PCR reaction primed with a pool of target-specific primers to enrich subsequent qPCR reactions when starting sample is limited.

**Table 1:**
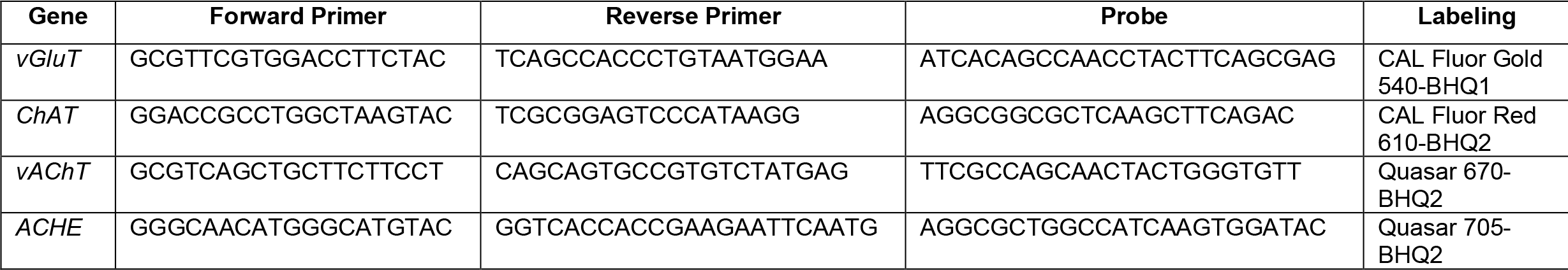
Forward and reverse primer pairs and associated probe sequences.

### Multiplex primer and probe design

Genes of interest were identified from the *C. borealis* nervous system transcriptome (Northcutt et al. 2016). Primer and probe sequences targeting *C. borealis* acetylcholinesterase (AChE), vesicular acetylcholine transporter (vAChT), choline acetyltransferase (ChAT), and vesicular glutamate transporter (vGluT) were designed using the RealTimeDesign™ qPCR Assay Design software (LGC Biosearch Technologies). Primer pairs and probes for the genes of interest were designed to be run in a single qPCR reaction. These assays relied on the following probe fluorophore/quencher pairs to detect PCR products produced during the qPCR reaction: (FAM-BHQ1, CAL Fluor Gold 540-BHQ1, CAL Fluor Red 610-BHQ2, Quasar 670-BHQ2 and Quasar 705-BHQ2). Forward and reverse primer pairs and associated probe sequences are reported in Table 1.

### Quantitative polymerase chain reaction (qPCR)

After preamplification, cDNA samples were diluted 7.5x in nuclease-free water (150 μL final volume). 10 μL qPCR reactions were carried out using PerfeCTa Multiplex qPCR ToughMix according to the manufacturer’s instructions (5X, QuantaBio). These reactions included pre-amplified cDNA (2.5 μL), forward and reverse primers (2.5 μM), and probes (0.3125 μM each probe). Reactions were run in triplicate on 96-well plates using a CFX96 Touch™ Real-Time PCR Detection System (Bio-Rad). Cycling conditions for qPCR reactions were as follows: 95°C for 3 min; 40 cycles of 95°C for 15 s and 58°C for 1 min. Fluorescent measurements were taken at the end of each cycle.

Conversion of quantitation cycle (C_q_) value to absolute copy number for each gene was estimated by interpolating C_q_ values for each gene into a standard curve of known copy number from 10^6^ to 10^1^ copies. We also accounted for the sample amount and the 14-cycle preamplification in our final estimation.

### Recording, Analysis and Statistics

Data were acquired using pClamp 9 (Molecular Devices), sampled at 5 kHz on a PC using a Digidata 1332A (Molecular Devices). Statistical and graphical analyses were done using GraphPad Prism 7.0 and Origin 8.5. For statistical analysis, mean, standard error of the mean and p-value are reported, unless indicated.

## Results

The electrically coupled AB and PD neurons comprise the pacemaker ensemble of the pyloric network. These neurons oscillate synchronously and together drive the rhythmic activity of the network by simultaneously inhibiting the pyloric follower neurons, including the LP and PY neurons. A cycle of the triphasic pyloric rhythm is generated by the synchronous burst of the pacemaker neurons, followed by a burst in the LP neuron and then a burst in the PY neurons (Fig 1A). Our goal is to confirm that, in *Cancer borealis*, the AB and PD neurons use different neurotransmitters (ACh and glutamate), and to identify the relative contributions of the AB and PD neurons to the composite synapses in the follower neurons LP and PY.

**Figure 1:**
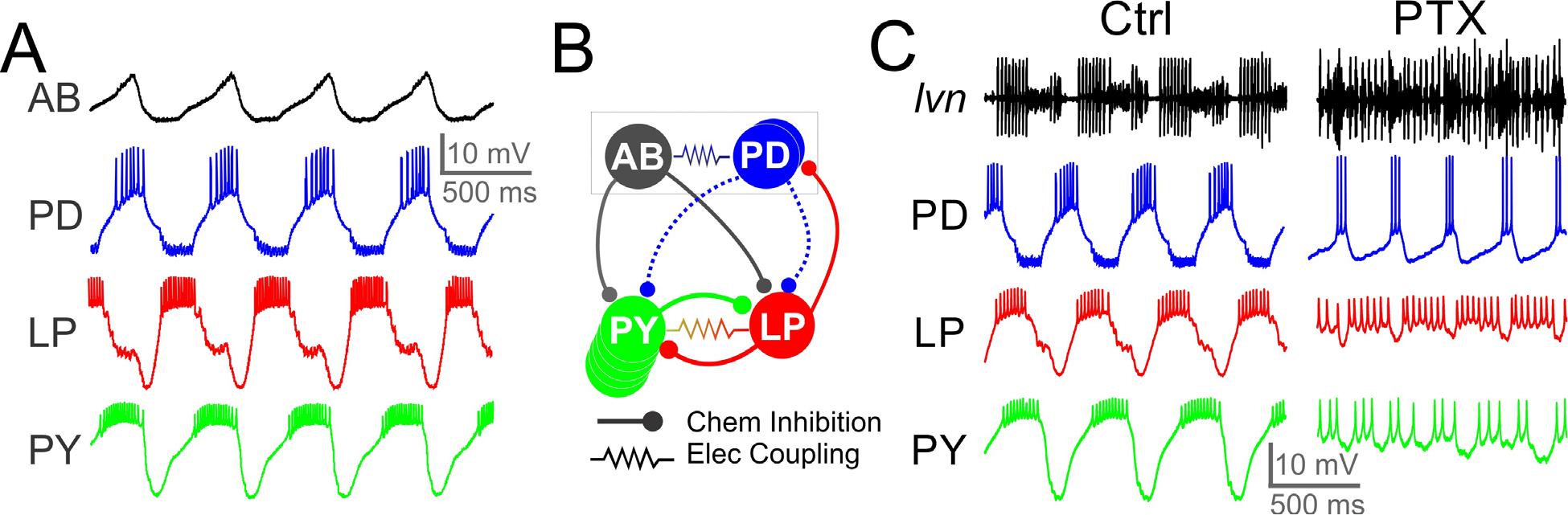
The Jonah crab pyloric circuit activity, connectivity and response to pharmacological block of glutamatergic synapses. **A.** Intracellular recordings of the AB, PD, LP and PY neurons during ongoing pyloric activity. **B.** Schematic diagram of the pyloric circuit, consisting of the AB and PD pacemaker ensemble neurons, and the follower LP and PY neurons. Putative cholinergic synapses from the PD neuron are shown as dashed curves. **C.** Simultaneous extracellular recording from the lateral ventricular nerve (*lvn*; containing the axons of PD, LP and PY neurons) and intracellular recordings of the PD, LP and PY neurons in control saline (Ctrl) and in the presence of 10^−5^ M picrotoxin (PTX) which blocks glutamatergic transmission in the STG. Rhythmic activity in PD continues in PTX, but the LP-induced IPSPs disappear. Residual rhythmicity remains in the LP and PYneurons.

### Single-cell expression of neurotransmitter genes from STG neurons

We first determined expression of mRNA levels for genes associated with neurotransmitter phenotype in single neurons: (1) acetylcholinesterase (AChE), (2) vesicular glutamate transporter (vGluT), (3) choline acetyltransferase (ChAT) and (4) vesicular acetylcholine transporter (vAChT). In these experiments, individual cells were identified and then assayed for gene expression (see Methods). Figure 2 shows the mean level of expression across the four neurons of interest. To determine if the neurons were cholinergic, we measured two markers for ACh expression: ChAT and vAChT. ChAT is responsible for the synthesis of acetylcholine, while vAChT is responsible for the transport of ACh from the cytoplasm into synaptic vesicles. Generally, neurons that express these two genes are considered to release ACh (Arvidsson et al. 1997). The PD neuron showed significantly higher expression of the two ACh markers when compared to other pyloric neurons (ChAT: Fig 2A; p<0.05, F(_3,17_)= 3.127, One-Way ANOVA, vAChT Fig 2B; p<0.001, F(_3,19_)=64.13, One-Way ANOVA). AB, LP, and PY neurons were effectively negative for detection of ChAT and vAChT expression.

**Figure 2:**
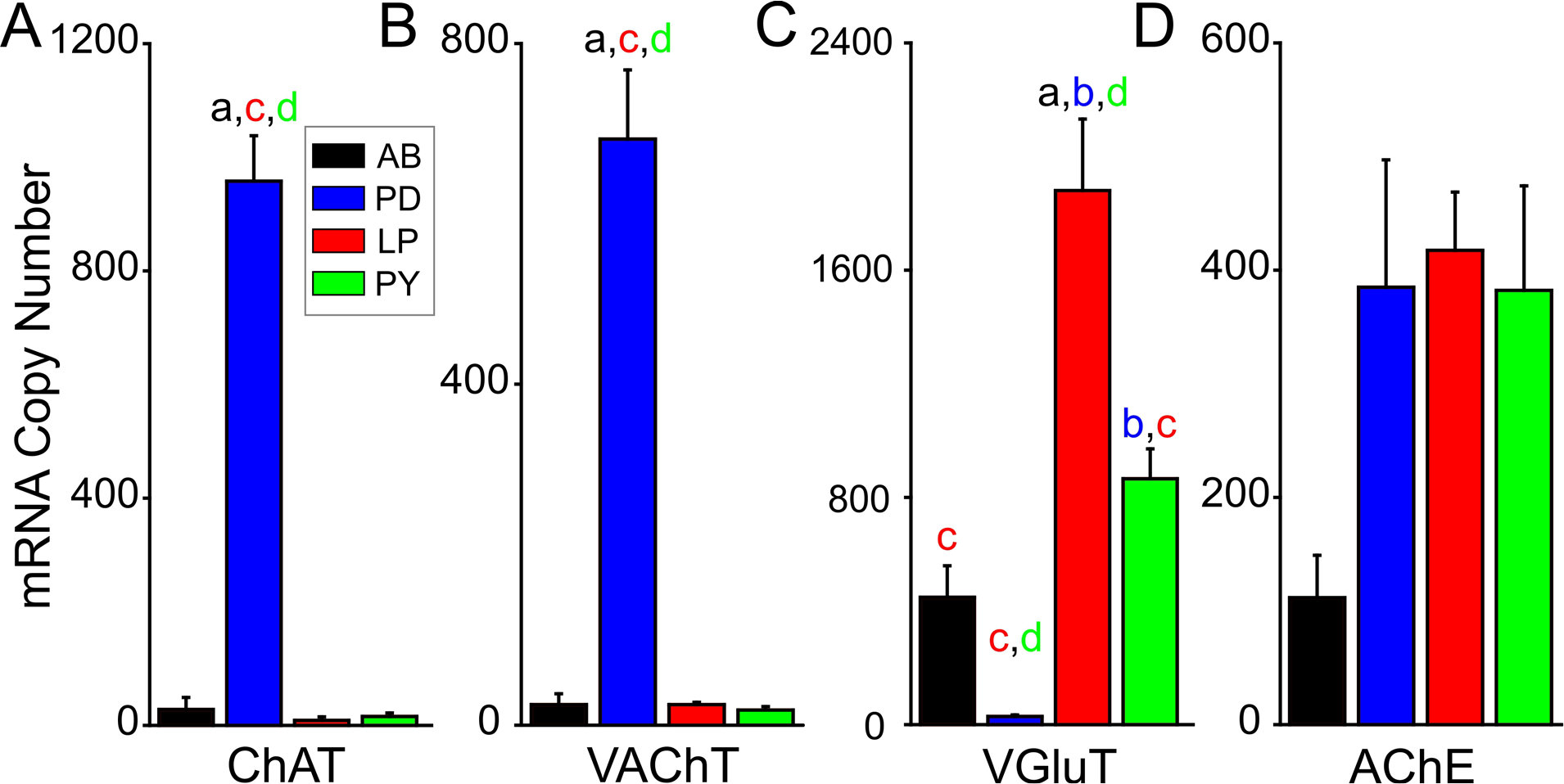
mRNA copy number for glutamatergic and cholinergic neurotransmission differs among pyloric circuit neurons in the Jonah crab. **A**. mRNA copy numbers for choline acetyltransferase (ChAT). The AB, LP and PY neurons show little expression, but the PD neuron has higher ChAT copy numbers (p<0.05). **B**. mRNA copy number for vesicular acetylcholine transporter (VAChT). As with ChAT, he PD neuron has the largest copy number for VAChT (p<0.0001). **C**. mRNA copy number for glutamate vesicular transporter (vGluT). The LP and PY neurons have large number of vGluT mRNA, however, the PD neuron expresses little vGluT mRNA compared to the other neuron types (p<0.0001). **D.** mRNA copy number for acetylcholinesterase (ACHE) is not significantly different among the four neuron types (p=0.066). For all measurements: AB (a) n=7; PD(b) n=5; LP (c) n=5; PY (d), n=6). Groups labeled with (a-d) indicate a significant difference (p<0.05) from one another through post-hoc analysis.

To determine glutamate expression across pyloric neurons we measured vGluT. vGluT is responsible for the transport of glutamate from the cytoplasm into synaptic vesicles, and is commonly used a marker of neurons that use glutamate as a neurotransmitter (Jing et al. 2015; Kolodziejczyk et al. 2008). PD neurons showed minimal expression for vGluT while all three other neurons showed significantly greater expression (Fig 2C; p<0.001, F(_3,19_)=19.27 One-Way ANOVA). This result strongly suggests that in *C. borealis*, like *P. interruptus*, the two pacemaker neurons PD and AB express different neurotransmitters, and that each pacemaker neuron may differentially inhibit the follower neurons. Lastly, the four neurons did not differ significantly in AChE expression (Fig 2D; p=0.066, F=2.86; One-Way ANOVA), which suggests that all four neuron types receive cholinergic synaptic input.

### Examining the PD-evoked LP, and PY neuron IPSP amplitudes

To measure the synaptic strength, we first examined the differences in amplitude of the pacemaker-evoked IPSPs in the LP and PY neurons in response to the PD neuron depolarization. The IPSP amplitude elicited in both LP and PY neurons increased with increased PD depolarization as expected from a graded synapse (Figs. 3A and 3B). To measure the synaptic activation curve, in each experiment and for reach of the two synapses, we fit the peak synaptic currents at all presynaptic membrane potentials with a Boltzmann sigmoidal curve (Fig 3B; see Methods) to estimate the PSP maximum amplitude (max IPSP), the half-activation voltage (V_mid_) and the slope factor (V_C_). The synaptic potential measured in the LP neuron was 36% larger in amplitude than that in the PY neuron (Student’s t test; p=0.033; N=12). However, there was no difference in the V_mid_ or V_C_ values (V_mid_: Student’s t test; p=0.221; N=12; V_C_: Student’s t test; p=0.797; N=12).

**Figure 3:**
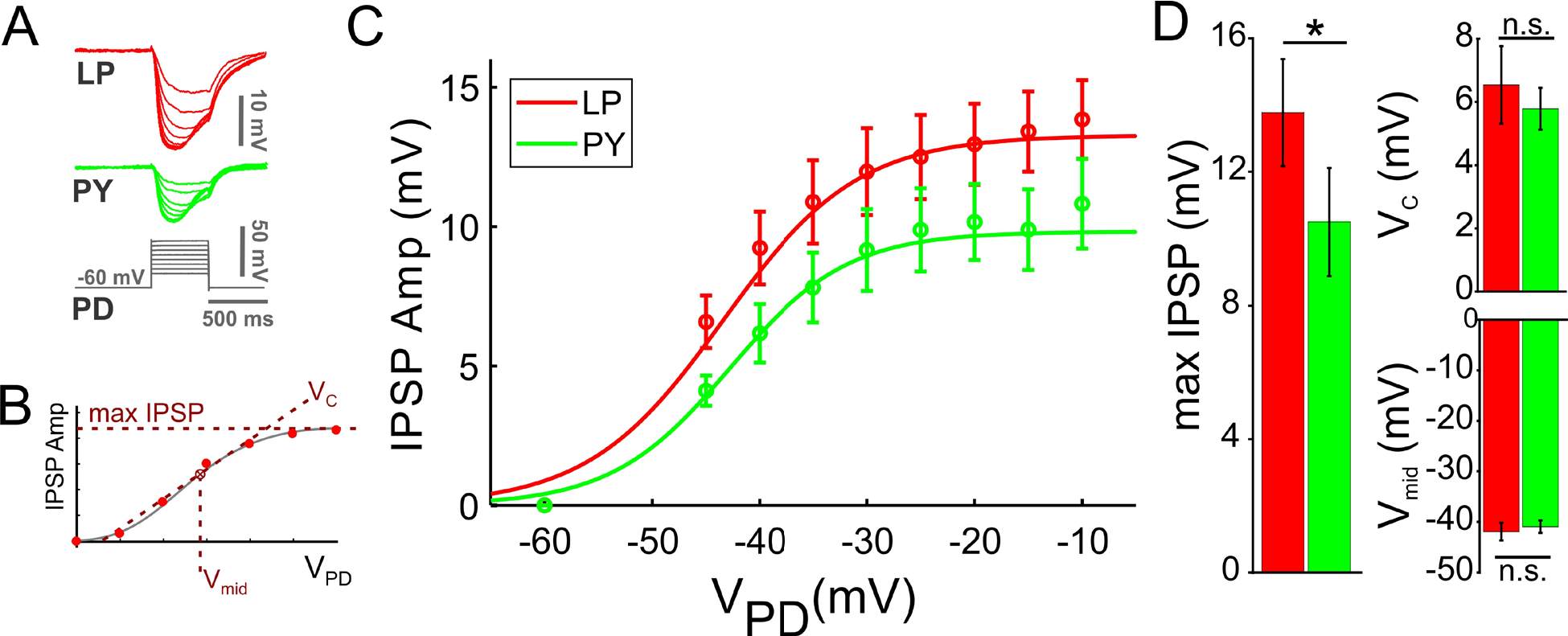
The pacemaker induced graded inhibitory postsynaptic potential is different in the follower PD and PY neurons. **A.** Voltage trace of the PD depolarization, and the simultaneously recorded IPSPs in the LP and PY neurons. **B.** To estimate the IPSP maximum amplitude (max IPSP), the half-activation voltage (V_mid_) and the slope factor (V_C_), in each experiment, we fit both the LP and PY peak IPSP amplitudes at all PD presynaptic membrane potentials with a Boltzmann sigmoidal curve. **C.** With increased PD depolarization, both the LP (red) and PY (green) IPSP increased in amplitude as expected in a graded synapse. **D.** The max IPSP for the LP (red) neuron was 36% larger than in the PY (green) neuron (p=0.033). There was no difference in the V_mid_ (p=0.221) or V_C_ values (p=0.797). N=12 experiments.

### Does the PD neuron have a functional synapse onto the LP and PY neurons?

The AB and PD neurons make a compound inhibitory synapse to both LP and PY neurons in the spiny lobster *P. interruptus* which consists of a comparable contribution from each of the two presynaptic neurons (Rabbah and Nadim 2007). Our results showed that in the crab *C. borealis*, the AB and PD neurons have different mRNA expressions for glutamate and acetylcholine (Fig 2). We therefore examined the relative contributions of the AB and PD neurons to the total synaptic effect in the LP and PY neuron in *C. borealis*.

We used picrotoxin (PTX, 10^−5^ M) to block the glutamatergic synapses from the AB neuron and measured the IPSPs using the presynaptic pulse protocol used above. In both synapses, PTX drastically reduced the amplitude of the IPSP, but did not completely block it (Fig 4A). For IPSPs measured in both the LP and the PY neurons, this effect was consistent at all presynaptic potentials (Fig 4B). A comparison of the synaptic activation curves in control and PTX showed that PTX significantly decreased the max IPSP amplitude in both synapses (LP: F(_2,18_)=18.47, p<0.0001, N=12, one-way RM-ANOVA; PY: F(_2,27_)=16.51, p<0.0001, N=11, one-way RM-ANOVA). The strength of the pacemaker induced synapse in the LP neuron in PTX was 24% of the control value, whereas in the PY neuron it was 29% of control. The effect of PTX on both synapses was partially washable (Fig 4). There was no difference in the V_mid_ or V_C_ values in Ctrl and PTX for both the LP and PY neurons (V_mid_: LP, p=0.115, PY, p=0.068; V_C_: LP, p=0.989, PY, p=0.464; one-way RM-ANOVA).

**Figure 4:**
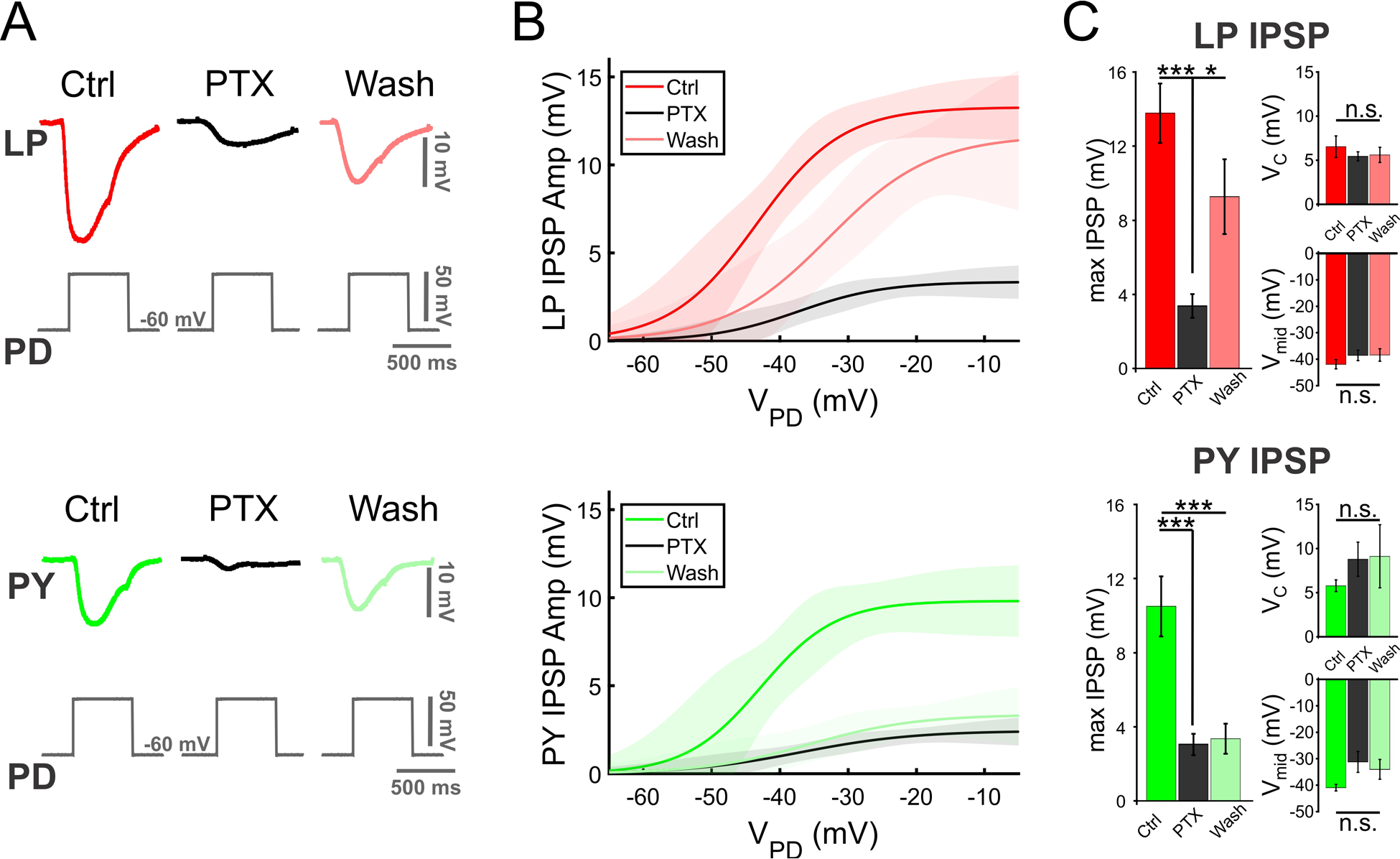
Blocking glutamatergic synapses blocks most but not all of the pacemaker induced postsynaptic potential in the follower neurons. **A.** Voltage trace of the PD depolarization with a 50 mV pulse, and the recorded IPSPs in the LP and PY neurons control, 10^−5^M picrotoxin (PTX), and wash. PTX reduced the amplitude of the IPSP in both LP and PY but did not eliminate it. **B.** Synaptic activation curve of the IPSP in the LP and PY neurons the three conditions. In both neurons, PTX reduced the IPSP amplitude but did not eliminate it. Activation curves are averages of fits to individual experiments as described in Fig 3B. Shaded area shows the 95% confidence interval. **C.** The strength of the pacemaker induced synapse was reduced significantly in PTX was 24% of the control value, and in the PY neuron it was (LP: 24% of control, p<0.0001; PY: 29% of control, p<0.0001). There was no difference in the V_mid_ (LP: p=0.115, PY: p=0.068) or V_C_ (LP: p=0.989, PY: p=0.464) values between Ctrl and PTX for both the LP and PY neurons. (***: p<0.0001; *: p<0.05.)

### Characterization of Synaptic Dynamics

To characterize the extent of short-term dynamics of the pacemaker synaptic input to follower LP and PY neurons, we depolarized the PD neuron to different membrane potentials with consecutive square pulses (five 500 ms pulses with 500 ms inter-pulse intervals; see Methods) and recorded the IPSPs in the LP and PY neurons (Fig 5A). The second and subsequent IPSPs in both the PY and LP neurons were smaller in amplitude than those elicited by the first pulse, indicating that both synapses had short-term depression. The extent of synaptic depression was quantified as a ratio of the amplitudes of the fifth (steady state) and first IPSPs (5^th^/1^st^) for each neuron (Mamiya et al. 2003). Although there was some variation in this paired-pulse ratio, the level of depression was statistically independent of the presynaptic voltage amplitude or the postsynaptic cell type (LP and PY, p = 0.9095, N=12, two-way ANOVA; Fig 5B).

**Figure 5:**
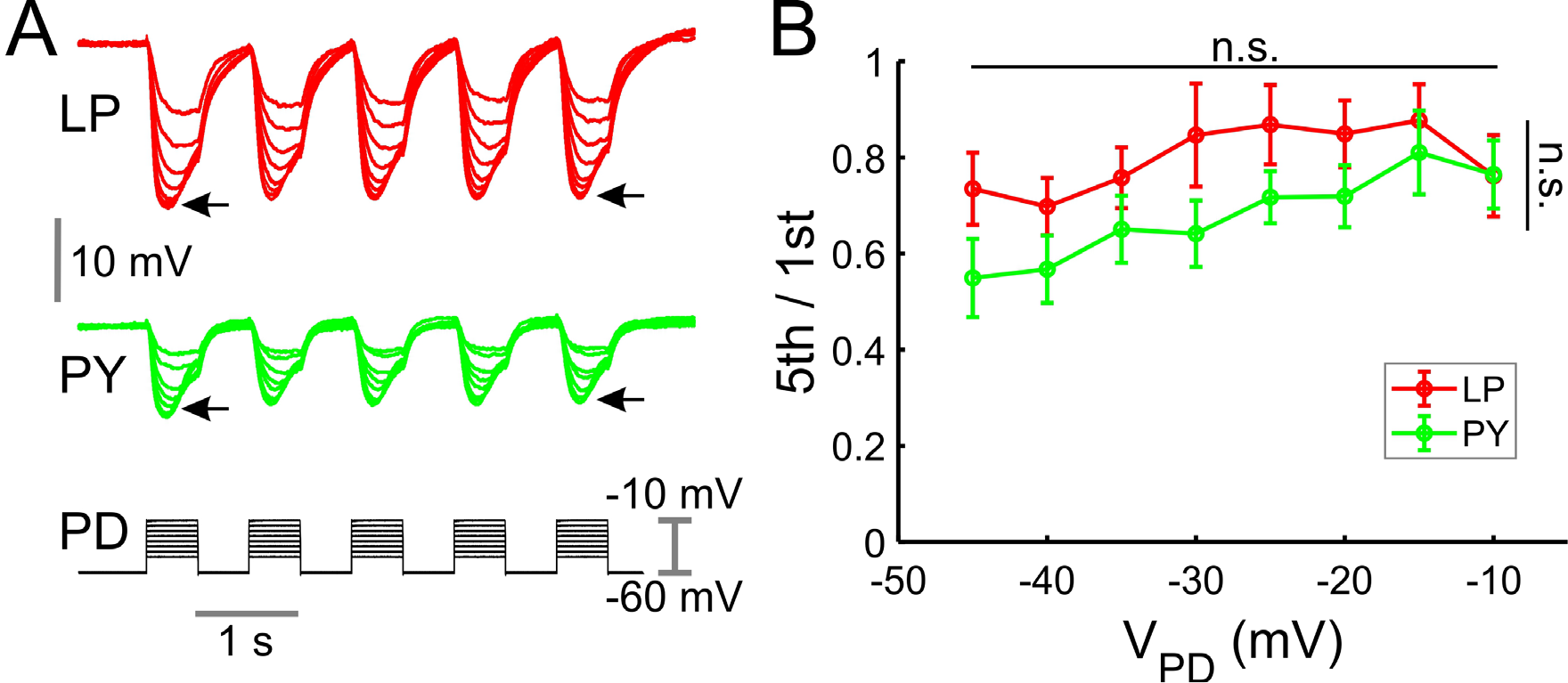
The pacemaker induced postsynaptic potentials in the LP and PY neuron show similar short-term depression. **A.** The PD neuron was depolarized with a series of five consecutive square pulses of different amplitudes, and the IPSPs in both the PY and LP neurons were recorded. The extent of the synaptic depression was quantified by taking the ratio of the amplitude of the fifth to first IPSP (5^th^/1^st^, denoted with arrows). **B.** Across all voltages measured, the level of depression (5^th^/1^st^ ratio) was independent of the presynaptic voltage or the postsynaptic neuron type (p = 0.9095).

To examine the contribution of the synaptic output of the PD neuron to synaptic depression, we compared the level of depression in the LP and PY neuron IPSPs in control and PTX saline (Fig 6A). Because presynaptic voltage amplitude did not influence depression levels, we only show the results for the 40 mV amplitude pulses. Once again, even though there was some variability in the paired-pulse ratio (5^th^/1^st^), we found no significant difference between depression levels in control, PTX or wash in either the LP or PY neuron IPSPs (LP: p=0.066, N=12; PY: p=0.973, N=11, one-way RM-ANOVA, Fig 6B). This result indicates that the level of depression is not significantly different for the synaptic output from the AB or PD neuron to these follower neurons.

**Figure 6:**
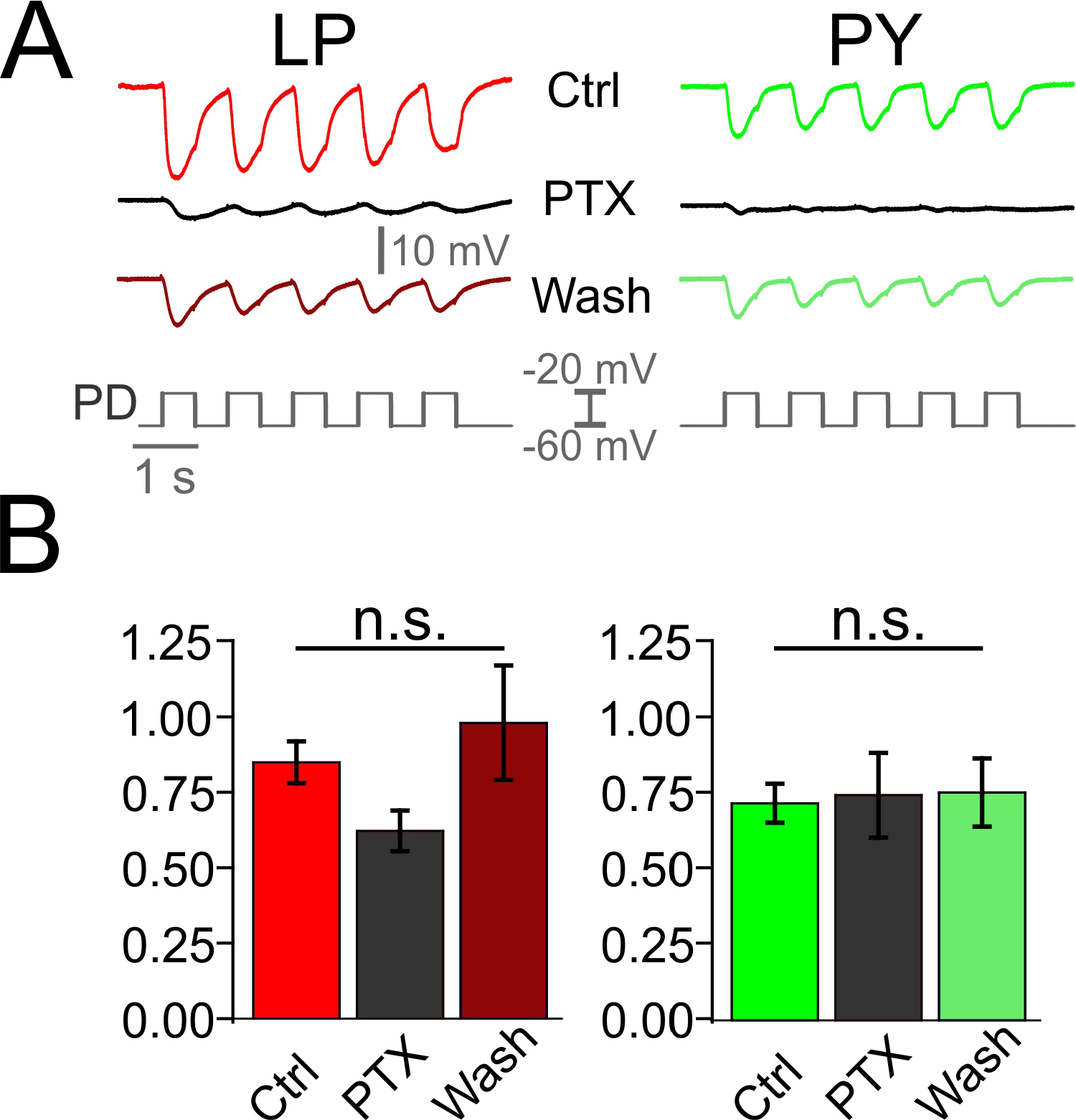
Short-term plasticity of the pacemaker induced synapses is independent of blocking glutamatergic synapses. **A.** The IPSPs recorded in the LP neuron and PY neurons, in response to five consecutive 40 mV square pulse depolarizations in the PD neuron, in control, PTX and Wash. **B.** We found no significant difference between depression levels (5^th^/1^st^ ratio) in control, PTX or wash in either the LP or PY neuron IPSPs (LP: p=0.066, PY: p=0.973).

## Discussion

Synaptic interactions are primary determinants of neuronal activity and circuit output (Nadim and Bucher 2014). Neurons that receive heterogeneous inputs naturally respond to a compound synaptic influence, which could be shaped by both the activity times of the presynaptic neurons and the properties of their synapses. In oscillatory networks, synaptic input arriving from synchronously active presynaptic neurons provide an instructive example of the direct influence of distinct synapses on the total compound synapse. A clear example of this is seen in pacemaker-driven oscillations involving a heterogenous group of pacemaker neurons (Rabbah and Nadim 2007).

Previous work in the spiny lobster *Panulirus interruptus* pyloric network demonstrated that the pacemaker group AB and PD neurons use different neurotransmitters to inhibit their circuit targets: the glutamatergic AB neuron produces fast inhibition, whereas the cholinergic PD neuron provides a delayed inhibition (Eisen and Marder 1982). In recent years, the Jonah crab *Cancer borealis* has become a primary species for studying properties of the stomatogastric nervous system. This shift has been due to numerous technical advantages of this species as a model system, as well as a number of seminal discoveries, including the identification and characterization of several modulatory projection neurons (Blitz and Nusbaum 2011; Nusbaum et al. 2001), the demonstration of ionic current expression variability and correlations (Khorkova and Golowasch 2007; Schulz et al. 2006; Temporal et al. 2012), the characterization of rules of peptide neuromodulation (Gray et al. 2017; Swensen and Marder 2001; 2000) and co-modulation (Li et al. 2018) and the development of molecular techniques (Northcutt et al. 2016; Schulz et al. 2007). Surprisingly, several basic circuit properties in the stomatogastric nervous system of this species remain unexplored and are assumed to be like those discovered in related species such as the spiny lobster. Among these assumptions is that the pyloric PD neurons are cholinergic and that their contribution to the total synaptic output of the pyloric pacemaker group is as described in the spiny lobster. Considering the differences in the circuit connectivity observed between the two systems (Marder and Bucher 2007; Stein 2009), and the central role of the pacemaker synapses in the pyloric circuit, it was important to clarify if the findings from the spiny lobster pacemaker synapses apply to the Jonah crab.

Our data confirmed that, as in *P. interruptus*, in *C. borealis* the AB neuron is glutamatergic whereas the PD neuron is cholinergic. The differential use of neurotransmitters by the pacemaker neurons allows for the distinct regulation of the contribution of the AB or PD synapses to the follower neurons. For example, the combined pacemaker synapse to the LP and PY neurons has a relative contribution from AB and PD synapses that depends on the pacemaker cycle frequency and duty cycle. Additionally, the pyloric circuit activity is modified by endogenous neuromodulators, several which modulate the activity and synaptic output of the PD and AB neurons (Harris-Warrick and Johnson 2010; Li et al. 2018). In fact, in some cases, the AB and PD synapses are modulated differentially. For example, dopamine enhances the AB to LP synapse but almost abolishes the PD to LP synapse (Harris-Warrick et al. 1998). Consequently, the compound pacemaker synapse provides multiple degrees of flexibility for the influence of the pacemaker neurons on their targets, including most follower pyloric neurons.

The neurotransmitter phenotype of STG neurons was first described through enzyme activity assays (Marder 1976) and pharmacology of synaptic physiology (Marder and Eisen 1984a). These studies revealed that PD is cholinergic and AB, LP, and PY are likely glutamatergic. Since this early work, new molecular tools have made it possible to perform gene expression studies on multiple targets (Schulz et al. 2007) and even full transcriptomes in individual neurons (Cadwell et al. 2016). In the present work, we took advantage of single-cell quantitative PCR to measure the expression of mRNA that is commonly used to define neurons into cholinergic and glutamatergic phenotypes (e.g., Li et al. 1995; Roghani et al. 1994; Yamaguchi et al. 2007). We found that the PD neuron expresses high levels of ChAT and vAChT, but virtually no vGluT. This supports the original work using enzyme assays and synaptic pharmacology to identify the neurotransmitter phenotype in this neuron. Furthermore, we showed that AB, LP, and PY neurons express relatively high levels of vGluT, but almost undetectable quantities of ChAT and vAChT. These results are consistent with previous work that suggested, but did not definitively show, that these neurons are glutamatergic (Eisen and Marder 1982; Marder and Eisen 1984a). Given that PD neurons have now been shown to express mRNA consistent with cholinergic phenotype, have enzymatic activity for transporting acetylcholine, and have been shown physiologically to use acetylcholine as a neurotransmitter, our findings strongly suggest that mRNA expression of our candidate genes is, indeed, predictive of neurotransmitter phenotype in neurons of the STG. Therefore, we are confident that in this and future work can use these markers to define neurotransmitter phenotype in STG cells in the absence of direct synaptic or functional enzyme data.

To explore the functional strength of the AB and PD synapses to the follower neurons, we took advantage of picrotoxin, a known blocker of glutamatergic synapses in the stomatogastric ganglion (Bidaut 1980; Cleland and Selverston 1995; Marder and Paupardin-Tritsch 1978; Rinberg et al. 2013; Temporal et al. 2012). We found that, as in *P. interruptus*, in *C. borealis* the PD neuron makes a functional synapse onto the pyloric follower LP and PY neurons. The contribution of the PD neuron to the total compound synapse was about 24% in the LP neuron and about 29% in the PY neuron. This finding is consistent with the observation that, in PTX, the synapse from the pacemakers is often incapable of driving the follower neurons to produce a rhythmic oscillation (e.g., Fig 1, Rinberg et al. 2013).

The pyloric follower neurons LP and PY rebound from pacemaker inhibition to produce activity bursts at two different phases of each cycle, with the LP burst leading that of PY (see, e.g., Fig 1). It is natural to assume that this phase difference is partly due to different synaptic input strengths from the pacemaker neurons. However, previous studies of the *P. interruptus* pyloric circuit showed that this phase difference is entirely due to different intrinsic properties of these two follower neurons, and that the synaptic input from the pacemakers to these neurons is in fact identical (Rabbah and Nadim 2007; 2005). Our results comparing the corresponding synapses in *C. borealis* shows that in this species the pacemaker synaptic drive of the LP neuron is in fact stronger than that of PY. This is somewhat unexpected, if a weaker synaptic inhibition would allow for an earlier rebound, which is the opposite of what is seen from the LP and PY neuron burst phases. However, synaptic inhibition interacts with numerous intrinsic ionic current dynamics to produce post-inhibitory rebound. These include, the transient potassium current *I_A_*, the h current and low-threshold calcium currents such as the T current. It is plausible, for example, that smaller *I_h_* levels or larger *I_A_* in the PY neuron would compensate for the weaker inhibitory drive of this neuron to produce a delayed phase (Zhang et al. 2009). It is known that, in *P. interruptus*, both *I_A_* and *I_h_* levels in these neurons are modified by the same monoaminergic neuromodulators that also modulate the synaptic output from the AB and PD neurons (Harris-Warrick and Johnson 2010; Parker et al. 2018). It is therefore plausible that, in modulated states, the synaptic input from the pacemakers may have different relative strengths and different interactions with the intrinsic voltage-gated currents, thus leading to shifts in the relative phases of the follower LP and PY neurons.

In oscillatory circuits, short-term dynamics facilitate changes in synaptic strength as a function of network frequency (Abbott et al. 1997; Rothman et al. 2009). As in previous studies, our results show that the pacemaker synapses to these follower neurons showed depression (Li et al. 2018; Tseng et al. 2014). We also demonstrated that the level of depression is independent of the presynaptic voltage amplitude, the follower neuron type and the presence of PTX. Thus, both AB and PD synaptic outputs have similar short-term dynamics, which may indicate a coordinated adjustment of the synaptic strength as a function of network frequency.

Pacemaker neurons are involved in rhythm generation in a variety of networks (Del Negro et al. 2018; Grace 2016; Jarabo and Martin 2017; Koshiya and Smith 1999; Snider et al. 2018). In most networks, these neurons consist of a heterogeneous population that have multiple co-transmitters or, even in those with the same neurotransmitter identity, have distinct synaptic dynamics. Because these pacemaker neurons can vary in their synaptic release dynamics or be differentially targeted by neuromodulators, their synaptic influence is subject to the similar levels flexibility as that seen in the pyloric pacemaker neurons.

## Conclusions

Our work builds on previous studies of the pyloric network in other species of decapod crustaceans that reported on the neurotransmitter content, ion selectivity, and the short-term dynamics of the pacemaker to follower neuron synapses (Eisen and Marder 1982; Harris-Warrick and Johnson 2010; Li et al. 2018; Marder and Eisen 1984b; Nadim and Manor 2000; Tseng et al. 2014). The heterogeneity in neurotransmitter type and release properties of the Jonah crab pyloric pacemaker neurons provides this circuit with mechanisms to differentially control the relative contribution of each synapse. A systematic analysis of the differential neuromodulation and analysis of the synaptic dynamics is needed to provide a fuller understanding of the differential roles of the two pacemaker synapses in the biological context. Furthermore, a similar analysis could be applied to understand the synaptic properties of disparate synchronized input in other oscillatory to unravel the contribution of synaptic dynamics and its neuromodulation to circuit output.

